# Under-ice mesocosms reveal the primacy of light but the importance of zooplankton in winter phytoplankton dynamics

**DOI:** 10.1101/802819

**Authors:** Allison R. Hrycik, Jason D. Stockwell

**Affiliations:** University of Vermont, Rubenstein Ecosystem Science Laboratory, 3 College Street, Burlington, Vermont, USA 05401, ORCID: 0000-0002-0870-3398; University of Vermont Biology Department, 120A Marsh Life Science, 109 Carrigan Drive, Burlington, Vermont 05405, ORCID: 0000-0003-3393-6799

**Keywords:** Winter limnology, food webs, phytoplankton, zooplankton, rotifers

## Abstract

Factors that regulate planktonic communities under lake ice may be vastly different than the open-water season. However, under-ice food webs in temperate lakes are poorly understood, despite expected changes in light availability, ice cover, and snowfall associated with climate change. We hypothesized that light limitation (bottom-up control) outweighs zooplankton grazing (top-down control) on phytoplankton biovolume and community structure under ice in a north temperate lake. Using in situ under-ice mesocosms, we found that light had stronger effects on phytoplankton abundance than zooplankton, as expected. Specifically, low light limited growth of diatoms, cryptophytes, chrysophytes, and chlorophytes. Zooplankton, however, also significantly affected phytoplankton by decreasing diatoms and cryptophytes, in contrast to the common assumption that zooplankton grazing has negligible effects under ice. Ammonia and soluble reactive phosphorus decreased in high light treatments presumably through uptake by phytoplankton, whereas ammonia and soluble reactive phosphorus increased in high zooplankton treatments, likely through excretion. In situ experimental studies are commonly applied to understand food web dynamics in open-water conditions, but are extremely rare under ice. Our results suggest that changes in the light environment under ice have significant, rapid effects on phytoplankton growth and community structure and that zooplankton may play a more active role in winter food webs than previously thought. Changes in snow and ice dynamics associated with climate change may alter the light environment in ice-covered systems and significantly influence community structure.

## Introduction

The relatively recent and rapid changes in winter conditions in temperate zones have led to declining ice cover in temperate lakes (Sharma et al. 2019) and altered snow cover and snowmelt dynamics (Musselman et al. 2017). However, the impact of such changes in winter conditions on lake food web dynamics under ice is poorly understood (Salonen et al. 2009; Sommer et al. 2012) because winter limnology has been under-studied compared to the open water season (Salonen et al. 2009; Hampton et al. 2015). Experimental studies crucial to understand lake processes during the open-water season (e.g., Schindler 1977) are missing during the ice-covered season, despite the potential for winter plankton community dynamics to set inoculum conditions for the open water season. For example, phytoplankton can reach bloom densities under ice (Katz et al. 2015) and support abundant zooplankton populations (Grosbois and Rautio 2018). A study of 101 lakes worldwide found that winter chlorophyll-*a* (a proxy for phytoplankton abundance) reached an average of 43% of summer chlorophyll-*a* (Hampton et al. 2017). Winter phytoplankton populations may also be important as inoculum for spring blooms (Feuchtmayr et al. 2012). To better understand if and how changes in snow and ice cover will affect biotic communities under ice and potentially into the open water season, we must first disentangle the drivers of food web dynamics under ice.

Light is widely accepted as the major driver of phytoplankton production under ice because it limits photosynthetic activity and can be highly variable depending on winter conditions (Salonen et al. 2009; Hampton et al. 2015), including day length, ice thickness, ice clarity, and especially snow cover (Bolsenga and Vanderploeg 1992). Physical factors such as light and temperature are considered the main drivers of winter phytoplankton biovolume in the Plankton Ecology Group (PEG) Model that describes planktonic community succession in temperate lakes (Sommer et al. 1986, 2012). Predictably, total phytoplankton production is highest when light transmission is highest during winter (Maeda and Ichimura 1973).

Additionally, community structure may be highly sensitive to changes in light levels because different taxa may be better equipped to deal with different light conditions during winter. For example, diatoms have been found at bloom densities in conditions with clear ice and minimal snowpack (Katz et al. 2015). Some phytoplankton taxa with adaptations that allow them to succeed during light-limited conditions, such as mixotrophic or mobile flagellated taxa, are often found in high proportions under ice (Özkundakci et al. 2016). To this end, we may expect a higher proportion of known mixotrophic taxa, such as chrysophytes (Sanders et al. 1990), when light is limited, and higher total phytoplankton biovolume with high light transmission.

Zooplankton can control phytoplankton biovolume and community structure during the open water season through grazing, including selective feeding on specific phytoplankton groups (Bergquist et al. 1985). However, less is known about top-down effects of zooplankton under ice, which makes interpretation and prediction of food web interactions under ice and entering the spring phytoplankton bloom difficult (Sommer et al. 2012). Zooplankton actively feed under the ice (Vanderploeg et al. 1992; Grosbois and Rautio 2018), although they may be heavily dependent on accumulated lipid stores (Grosbois et al. 2017). Similar to the open water season, zooplankton grazing rates and phytoplankton response during winter are expected to be dependent on the zooplankton and phytoplankton species present, and their interactions. For example, winter zooplankton communities are often dominated by copepods and rotifers (Blank et al. 2009). A higher ratio of herbivorous to predatory rotifers may be expected under ice if *Daphnia* are limited (Obertegger et al. 2011). Copepods may have particularly strong impacts on phytoplankton community structure through selective raptorial feeding (Sommer et al. 2003), suggesting that winter zooplankton communities that are actively feeding may influence under-ice phytoplankton community structure and biovolume.

Changes in nutrient concentrations under ice may be closely linked to changes in phytoplankton and zooplankton communities. Nutrients are generally not expected to limit phytoplankton growth during winter, especially in eutrophic systems (Sommer et al. 2012). However, we may still expect a signal from phytoplankton communities in nutrient concentrations. Higher phytoplankton growth generally corresponds with reductions in forms of nitrogen and phosphorus that can be assimilated quickly, such as soluble reactive phosphorus (SRP), ammonia, and sometimes nitrate (Glibert et al. 2016). Manipulation of zooplankton biomass may also alter nutrient levels through excretion (Oliver et al. 2015). Other sources of nutrient inputs that are important in the open-water season, including phosphorus release from sediment (Penn et al. 2000), may also be a significant source of phosphorus under ice if oxygen is limited (Joung et al. 2017). We expected that nutrient concentrations would respond to changes in plankton communities but not to the extent that they affected phytoplankton growth (Sommer et al. 2012).

In this study, we used an in situ under ice carboy experiment to test the relative importance of zooplankton grazing versus light limitation on winter phytoplankton biovolume, community structure, and nutrient concentrations in a north temperate lake. We hypothesized that both low light and high zooplankton grazing would decrease phytoplankton biovolume, but that light would be quantitatively more important than zooplankton to drive total phytoplankton biovolume and have a larger impact on phytoplankton community structure under ice. Our hypothesis follows the PEG Model (Sommer et al. 1986) and its recent update (Sommer et al. 2012), in which physical factors are thought to shape winter phytoplankton communities compared to higher influence of zooplankton grazing and nutrient limitation during the open water season. To our knowledge, our under-ice carboy experiment is the first application of this type of mesocosm experiment under ice, despite wide application of carboy experiments during the open water season.

## Methods

The experiment took place in Shelburne Pond, Vermont, a small, hypereutrophic system with a mean depth of 3.4 m and maximum depth of 7.6 m (Vermont Department of Environmental Conservation 2018). We initiated the experiment over 2 consecutive days with 12 carboys on 25 January and 12 carboys on 26 January 2018. The transparent carboys were deployed on the north end of Shelburne Pond (44.39388°N, 73.16278°W) in an area with 0-2 cm of patchy snow on top of 30 cm of secondary ice (also called black ice; Block et al. 2018) and a water column depth of 4.6 m. We set carboys in a randomized grid pattern of six carboys by four carboys spaced 5 m apart. Each carboy was suspended by steel cable approximately 50 cm below the ice (Fig. 1A).

**Fig. 1.**
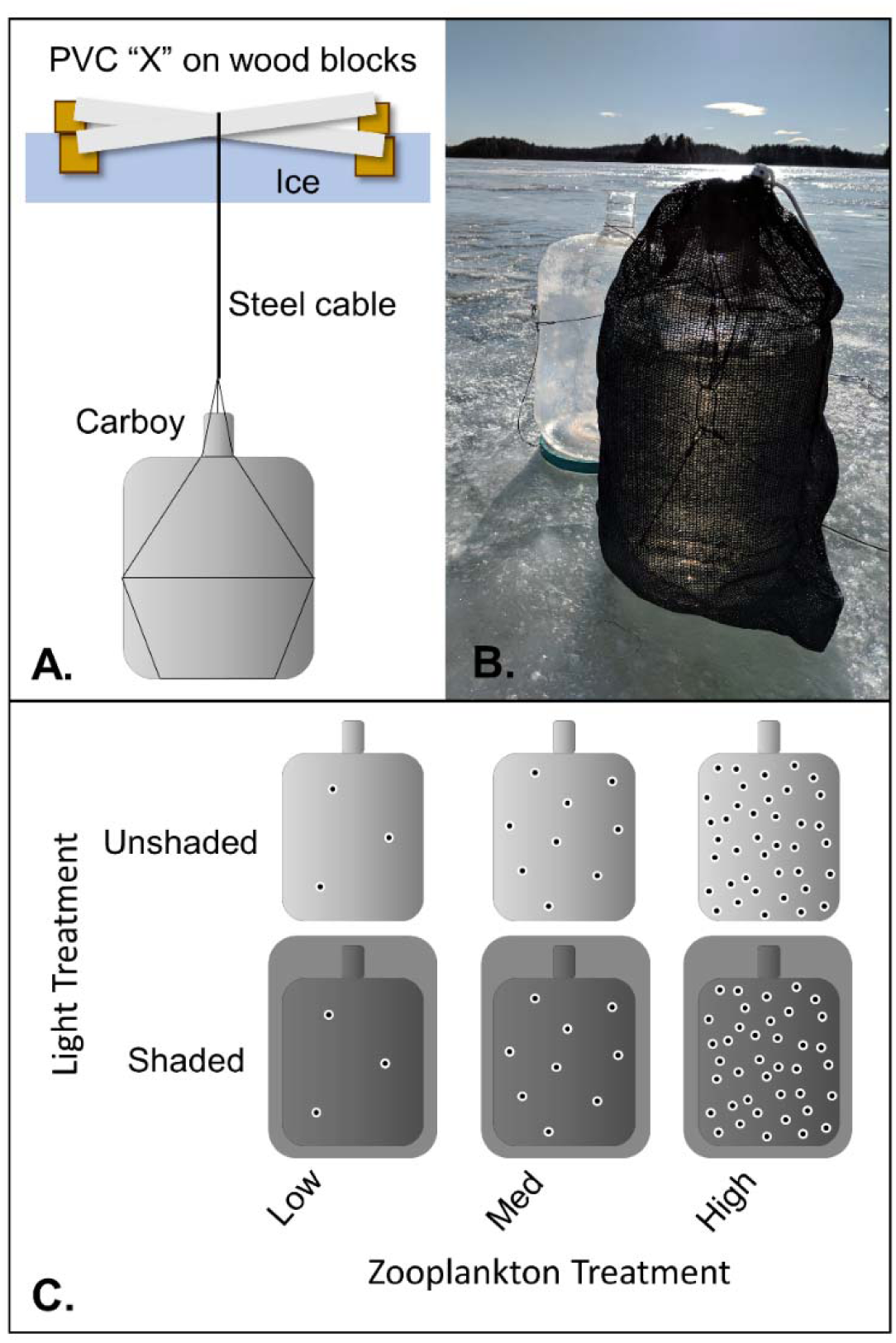
Carboy experimental design. A.) Each carboy was suspended in a randomized grid approximately 0.5 m below the ice by a steel cable harness connected to a PVC anchor in the shape of an “X.” PVC was placed on loose wood blocks to prevent it from freezing into the ice. B.) Greenhouse shade cloth covers blocked 85% of incoming light to simulate snow cover (photo credit: Hannah Lachance). C.) We crossed two light levels (unshaded and shaded) with three zooplankton levels with four replicates per treatment (see Results for zooplankton abundances).

We tested four replicates for each of six treatments: (1) low zooplankton/unshaded, (2) low zooplankton /shaded, (3) medium zooplankton/unshaded, (4) medium zooplankton/shaded, (5) high zooplankton/unshaded, and (6) high zooplankton/shaded (Fig. 1B/C). We mixed ambient water for all treatments in a 208-L plastic barrel. The barrel was filled by raising and lowering the intake of a hand pump throughout the top 4.0 m of the water column and filtering through a 350-µm sieve to remove large zooplankton. Pilot work indicated 350-µm was the best mesh size to remove large grazing zooplankton but kept most colonial phytoplankton. The sieved ambient lake water in the barrel was mixed constantly as each group of six 22.7-L carboys (i.e., one replicate for each treatment) was filled.

We then added zooplankton treatments directly to carboys. Two of the six carboys had no zooplankton added, two had ambient zooplankton added, and two had 10X natural abundance of large (i.e., sieved) zooplankton added. Zooplankton for the treatments were collected with a 13-cm diameter, 64-µm Wisconsin net through the upper 4.0 m of the water column and then retained on a 350-µm sieve. Desired densities were achieved for the ambient zooplankton treatment by using a plankton splitter and adding half of the sieved net haul to each of the two ambient abundance treatments because half of the volume strained for a 4-m tow was approximately equal to the carboy volume. The high zooplankton treatments had sieved zooplankton from five zooplankton net tows added to each carboy. Finally, we covered one carboy from each zooplankton treatment with greenhouse shade cloth that blocked 85% of incoming light to simulate the light-limiting effects of snow cover. The set-up process was repeated twice on 25 January and twice on 26 January for a total of 4 replicates per treatment. All setup processes, including filtering zooplankton, were performed in the field at the time of deployment. We affixed a MK-9 light and temperature sensor (Wildlife Computers Inc., Redmond, WA, USA) to the outside of one carboy without shade cloth and between another carboy and its shade cloth on 25 January. We also added a HOBO temperature sensor (Onset Computer Corporation, Bourne, MA, USA) to the inside of one shaded and one unshaded carboy on 26 January. Light values from MK-9 sensors were converted from relative units to µE·m^−2^·s^−1^ (Kotwicki et al. 2009). We did not leave any head space in the carboys to simulate the sealed conditions of an ice-covered lake. The ice over the carboys re-froze within one day of deployment.

Carboys were extracted from the lake 14 days after deployment. Although most summer carboy experiments are much shorter in duration (e.g., Griniene et al. 2016), we expected that slow phytoplankton growth rates at low temperatures (Cloern 1977) would necessitate a longer incubation time. Each carboy was inverted several times to homogenize contents before opening. We sampled phytoplankton, nutrients, and zooplankton from each carboy. We collected three phytoplankton samples by filling 125-mL bottles with water and its associated phytoplankton community, which was later measured out to 100 mL and preserved in Lugol’s solution. One 500-mL water sample was taken per carboy, and split into portions for different nutrient analyses once back at the lab as follows. A 100-mL sample was preserved with three drops of sulfuric acid to achieve a pH of 2.0 for later analysis of total phosphorus (TP), total nitrogen (TN), and total organic carbon (TOC). Two 45-mL samples were filtered through 0.45-µm syringe filters and frozen: one for soluble reactive phosphorus (SRP), and one for ammonia and nitrate + nitrite (NO_x_) quantification. We strained the remaining 21.84 L of water through a 20-µm Wisconsin net to sample crustacean zooplankton and rotifers. All zooplankton and rotifers were anesthetized with Alka Seltzer before preservation with 70% ethanol.

We identified phytoplankton to genus and counted full fields of view at 400x until reaching at least 300 natural units (cells for single-celled species or colonies for colonial species). We measured ten natural units per genus for each sample to calculate biovolume using Spot Basic software (Spot Imaging, Sterling Heights, MI). Dimensions measured were dependent on phytoplankton taxa present, e.g., we measured diameter for spherical cells and diameter and length for ellipsoid cells (Hillebrand et al. 1999). Within colonies, we measured ten individual cells per colony when possible. If fewer than ten cells were present or clearly visible, we measured all cells. We used those median measurements to calculate taxon-specific biovolume for each sample (Hillebrand et al. 1999), then taxon-specific biovolume was multiplied by cell abundance to estimate total biovolume per sample for each taxon. We only processed one out of the three phytoplankton samples collected per carboy because replicates within each carboy were very similar and would not have added to statistical power due to pseudoreplication. Analysis of three pairs of phytoplankton samples from the same carboys showed an average of 4.2% difference in cell counts for each genus.

We processed rotifers and crustacean zooplankton by measuring and counting at least 200 individuals per sample (200 rotifers and 200 zooplankton). Rotifers were counted and measured on a Nikon Eclipse Ni-U compound microscope (Nikon, Tokyo, Japan) with Spot Basic software (Spot Imaging, Sterling Heights, Michigan, USA), while zooplankton were identified and enumerated on an Olympus SZX12 dissecting microscope (Olympus Corporation, Tokyo, Japan) interfaced with a GTCO CalComp digitizer for measurements (Turning Technologies, Inc., Youngstown, Ohio, USA). Rotifer and crustacean zooplankton biomass were calculated using length-to-biomass conversions (all crustacean zooplankton and most rotifers) or length/width-to-biomass conversions (*Filinia* rotifers) (Watkins et al. 2011; United States Environmental Protection Agency 2016).

We measured nutrient concentrations primarily to ensure that we did not artificially limit nutrients in our study. Nutrient samples were either stored frozen (SRP, ammonia, and NO_x_) or acidified and refrigerated (TN, TOC, and TP) until analysis. We measured TN and TOC on a TOC-L total organic carbon analyzer with a TNM-L TN measuring unit (Shimadzu Corporation, Kyoto, Japan). We analyzed TP and SRP using the molybdenum colorimetry method (USEPA 1993) with ascorbic acid modification and a persulfate digestion for TP on a Shimadzu UV-VIS 2600 spectrophotometer (Shimadzu Corporation, Kyoto, Japan). Ammonia and NO_x_ were measured on a SEAL AA3 continuous flow autoanalyzer (SEAL Analytical, Mequon, Wisconsin, USA) using Method No. G-171-96 Rev. 15 with salicylate for ammonia and Method No. G-172-96 Rev. 18 for NO_x_.

All response variables (phytoplankton abundance and biovolume, rotifer abundance and biomass, and nutrient concentrations) were analyzed using 2-way ANOVAs (α=0.05) with zooplankton and light as predictor variables. We then used the ANOVA output for variance partitioning to quantify the contribution (partial *R*^*2*^) of zooplankton and light separately on response variables as well as the contribution of an interaction term between light and zooplankton. The values are reported as *R*^*2*^_*zoop*_, *R*^*2*^_*light*_, and *R*^*2*^ _*light:zoop*_.

We performed non-metric multidimensional scaling (nMDS) with Bray-Curtis distance on phytoplankton species composition to visualize differences in phytoplankton communities between treatments using the R package “vegan” (version 2.4.3; Oksanen et al. 2013). We removed the small number of phytoplankton records where phytoplankton could not be identified to genus, which should not significantly alter nMDS interpretation (Pos et al. 2014). We used perMANOVA with Bray-Curtis distance and 999 permutations to test significance of light and zooplankton treatments on overall phytoplankton community composition (Anderson 2001; Oksanen et al. 2013).

## Results

Light and temperature behaved as expected throughout the experiments. Light was reduced by greenhouse shade cloth (Supporting Information Fig. S1). Temperature remained consistent between light and shaded treatments, although internal carboy temperatures were slightly higher than external water temperatures. However, the difference between internal and external temperature (<0.3°C) was small compared to the overall increase in water temperature over the course of the experiment (Fig. S1).

Zooplankton abundance and biomass were significantly different between treatments (ANOVA; *F*_1,22_ = 225.1, *p* < 0.0001), but differences were not as great as intended (Fig. 2). Our low zooplankton treatments averaged (± SD) 64.5 ± 14.21 µg dry/L, medium zooplankton treatments averaged 99.8 ± 5.36 µg dry/L, and high zooplankton treatments averaged 338.6 ± 56.93 µg dry/L. That is, our intended “10x” zooplankton level had 3.4x the zooplankton biomass as our intended “1x” treatment. Consequently, we refer to zooplankton levels as low, medium, and high rather than 0x, 1x, and 10x. Zooplankton biomass was dominated by *Diacyclops thomasi* and zooplankton abundance was dominated by both *D. thomasi* and copepod nauplii (Fig. S2). Crustacean zooplankton body size followed a bimodal distribution with a smaller peak that represented copepod nauplii and *Chydorus spp.* and the larger peak represented adult zooplankton (Fig. S3). The smaller zooplankton were depleted in treatments with higher densities of large-bodied zooplankton (Fig. S3).

**Fig. 2.**
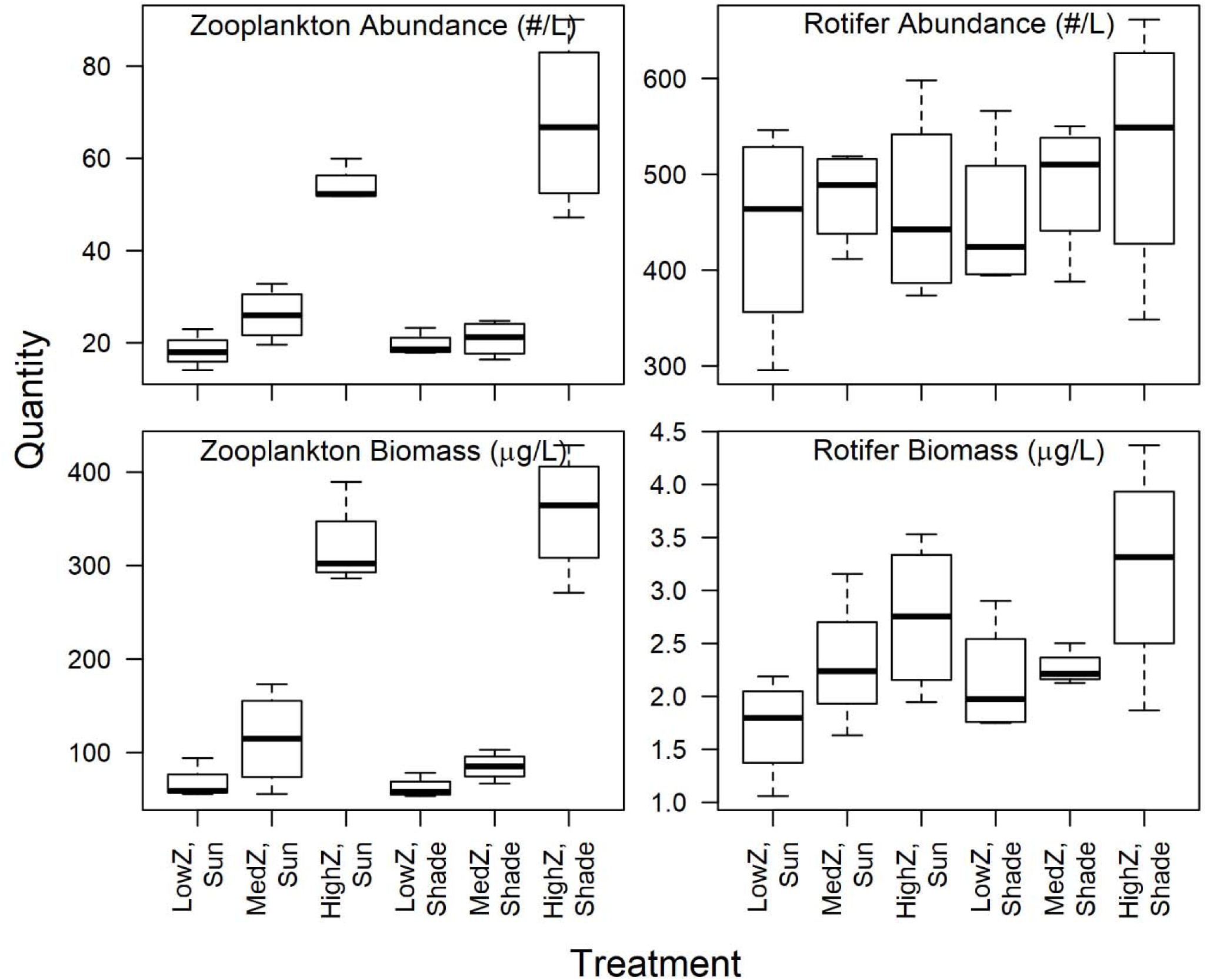
Crustacean zooplankton and rotifer abundance and biomass. “Z” indicates zooplankton level. Center lines indicate medians, boxes indicate first and third quartiles of data, and whiskers indicate minimum and maximum of data.

Differences in phytoplankton community composition among treatments was primarily driven by light, although some groups were also significantly affected by zooplankton levels. The biovolume of total phytoplankton and chlorophytes, and abundance and biovolume of diatoms, cryptomonads, and chrysophytes significantly increased with light (Table 1; Fig. 3; Fig. 4). Light most strongly increased diatoms (*R*^*2*^_*light*_ = 0.73 for abundance and *R*^*2*^_*light*_ = 0.62 for biovolume) and chrysophytes (*R*^*2*^_*light*_ = 0.76 for abundance and *R*^*2*^_*light*_ = 0.71 for biovolume) (Table 1). Zooplankton significantly decreased diatom abundance and cryptomonad biovolume but explained less variation overall than light treatments (*R*^*2*^ _*zoop*_ = 0.07 for diatom abundance and *R*^*2*^ _*zoop*_ = 0.33 for cryptophyte biovolume) (Table 1). Cyanobacteria and haptophytes were also found in all samples but did not vary by treatment (Table 1). Phytoplankton samples comprised diatoms, chlorophytes, cyanobacteria, cryptomonads, dinoflagellates, chrysophytes, haptophytes, synurophytes, desmids, and euglenoids (Table 2).

**Table 1.** 2-way ANOVA and variance partitioning results for each response variable. *N*=24 for each test (4 replicates per treatment). Dinoflagellate, desmid, euglenoid, and synurophyte phytoplankton abundance and biovolume were not analyzed statistically due to low sample size but are included in total phytoplankton calculations. Significant *p*-values (α=0.05) are bolded with their respective *R*^*2*^s.

| Response variable | $R^2_{light}$ | $R^2_{zoop}$ | $R^2_{light:zoop}$ | $p_{light}$ | $p_{zoop}$ | $p_{light:zoop}$ |
| --- | --- | --- | --- | --- | --- | --- |
| Total phytoplankton abundance | 0.07 | 0.08 | 0.01 | 0.2345 | 0.4637 | 0.8896 |
| Chlorophyte abundance | 0.00 | 0.02 | 0.06 | 0.7676 | 0.8317 | 0.5424 |
| Diatom abundance | **0.73** | **0.07** | 0.04 | **4.068\*10<sup>-8</sup>** | **0.0382** | 0.1643 |
| Cyanobacteria abundance | 0.00 | 0.04 | 0.03 | 0.7736 | 0.6524 | 0.7750 |
| Cryptomonad abundance | **0.37** | 0.12 | **0.16** | **0.0004** | 0.0659 | **0.0369** |
| Chrysophyte abundance | **0.76** | 0.05 | 0.02 | **5.429\*10<sup>-8</sup>** | 0.1128 | 0.4209 |
| Haptophyte abundance | 0.00 | 0.00 | 0.02 | 0.9256 | 0.9865 | 0.8169 |
| Total phytoplankton<br>biovolume | **0.51** | 0.10 | 0.01 | **0.0001** | 0.1366 | 0.8210 |
| Chlorophyte biovolume | **0.21** | 0.08 | 0.02 | **0.0306** | 0.3521 | 0.7309 |
| Diatom biovolume | **0.62** | 0.00 | 0.00 | **3.222\*10<sup>-5</sup>** | 0.9242 | 0.8959 |
| Cyanobacteria biovolume | 0.02 | 0.05 | 0.06 | 0.4892 | 0.5734 | 0.5450 |
| Cryptomonad biovolume | **0.23** | **0.33** | 0.01 | **0.0067** | **0.0064** | 0.8959 |
| Chrysophyte biovolume | **0.71** | 0.05 | 0.03 | **4.018\*10<sup>-7</sup>** | 0.1609 | 0.3427 |
| Haptophyte biovolume | 0.00 | 0.05 | 0.04 | 0.8360 | 0.6463 | 0.7056 |
| Rotifer abundance | 0.03 | 0.03 | 0.02 | 0.4581 | 0.4174 | 0.5253 |
| Rotifer biomass | 0.04 | **0.33** | 0.00 | 0.2867 | **0.0041** | 0.6723 |
| Total phosphorus | 0.11 | 0.02 | 0.03 | 0.1291 | 0.5490 | 0.4007 |
| Soluble reactive<br>phosphorus | **0.73** | **0.15** | 0.00 | **8.079\*10<sup>-10</sup>** | **9.489\*10<sup>-5</sup>** | 0.6969 |
| Total organic carbon | 0.00 | 0.00 | 0.01 | 0.7872 | 0.8323 | 0.6004 |
| Total nitrogen | 0.08 | **0.20** | 0.09 | 0.1350 | **0.0211** | 0.1080 |
| Ammonia | **0.72** | **0.21** | 0.00 | **2.342\*10<sup>-12</sup>** | **8.690\*10<sup>-8</sup>** | 0.9861 |
| Nitrate | 0.01 | 0.00 | 0.00 | 0.7133 | 0.8282 | 0.8996 |

**Table 2.**
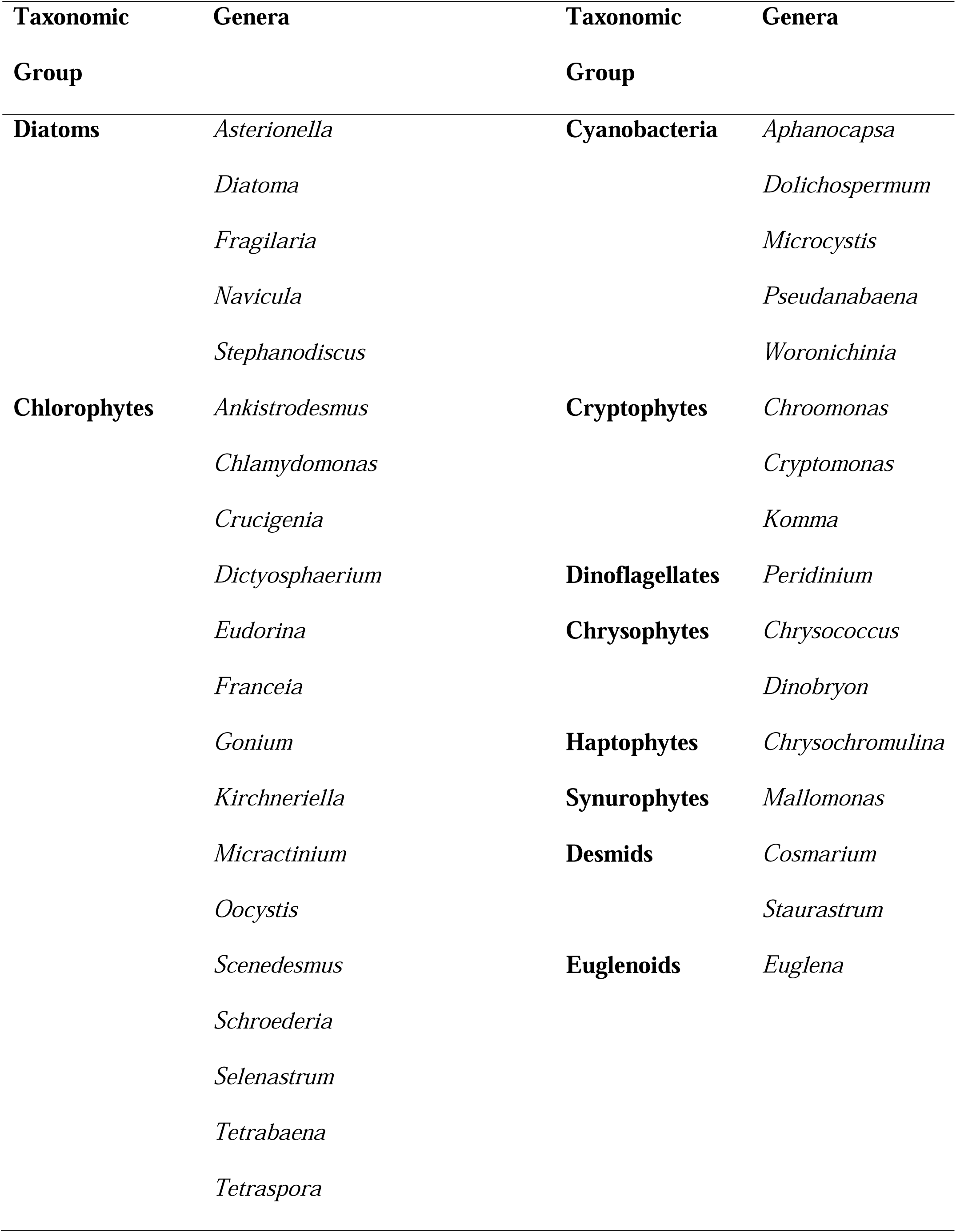
Phytoplankton genera found in mesocosms.

**Fig. 3.**
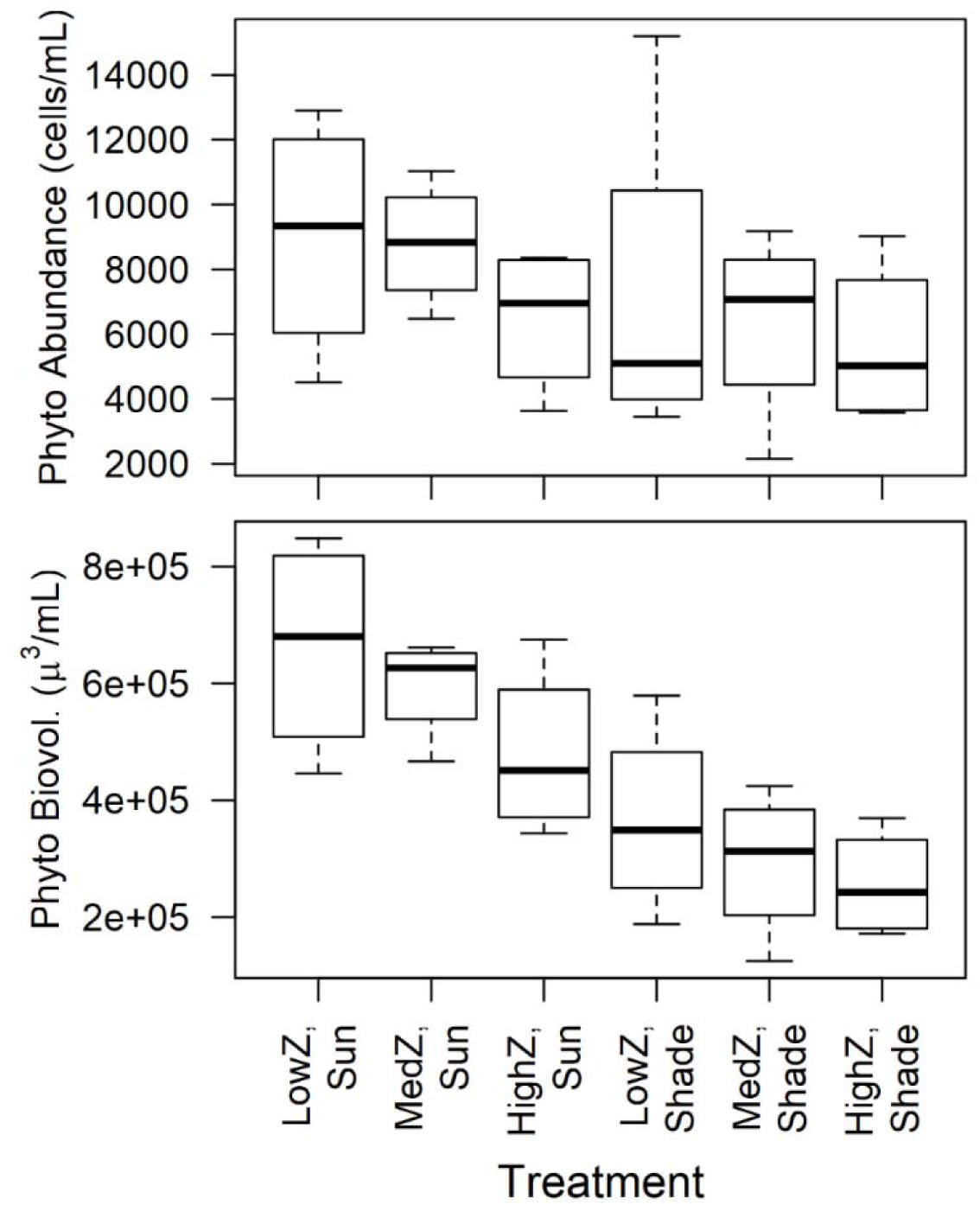
Total phytoplankton biovolume and abundance. “Z” indicates zooplankton level. Center lines indicate medians, boxes indicate first and third quartiles of data, and whiskers indicate minimum and maximum of data.

**Fig. 4.**
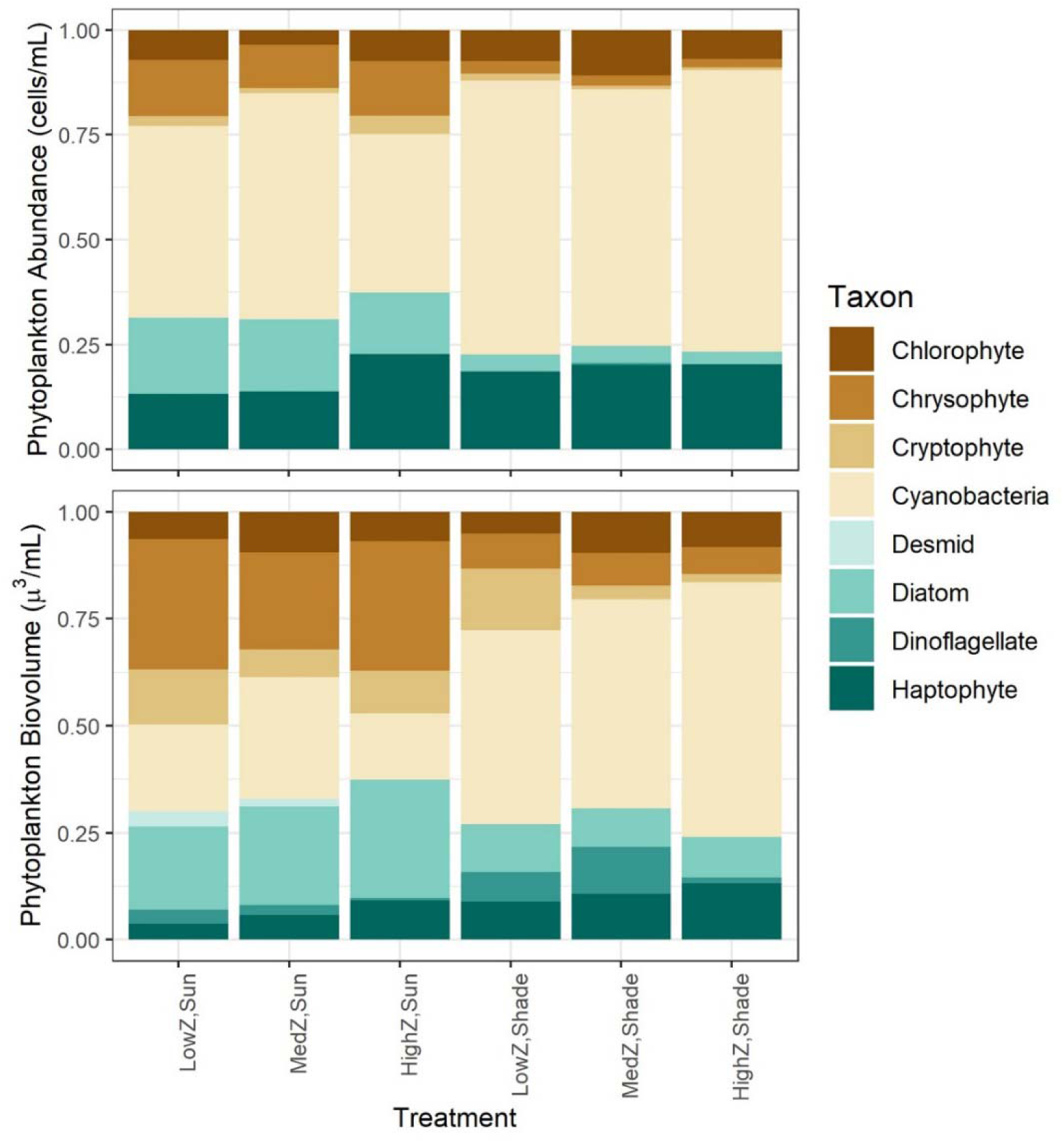
Phytoplankton proportional abundance and biovolume by taxonomic group. “Z” indicates zooplankton level.

Non-metric multidimensional scaling resulted in a clear distinction in phytoplankton communities between shaded and unshaded treatments for all axes, but little effect from zooplankton (Fig. 5). Treatments at the beginning of experiments were more similar to shaded treatments than unshaded treatments. Our final ordination had three axes to reduce stress from 0.22 with two axes to 0.15 with three axes (Fig. 5). Phytoplankton genera that drove separations along nMDS axes were mostly rare species that were only found in some treatments such as *Crucigenia, Diatoma, Aphanocapsa*, and *Dictyosphaerium* for the first axis and *Oocystis, Stephanodiscus, Selenastrum*, and *Gonium* for the second axis (Fig. S4). More common genera such as *Chrysochromulina* and *Woronichinia* were found across treatments, so contributed little to separations in nMDS axes (Fig. S4). PerMANOVA indicated a significant effect of light treatment (*pseudoF*_1,20_ = 12.3; *p*=0.001) but no effect of zooplankton treatment (*pseudoF*_2,20_ = 1.36; *p*=0.184) on overall phytoplankton community composition.

**Fig. 5.**
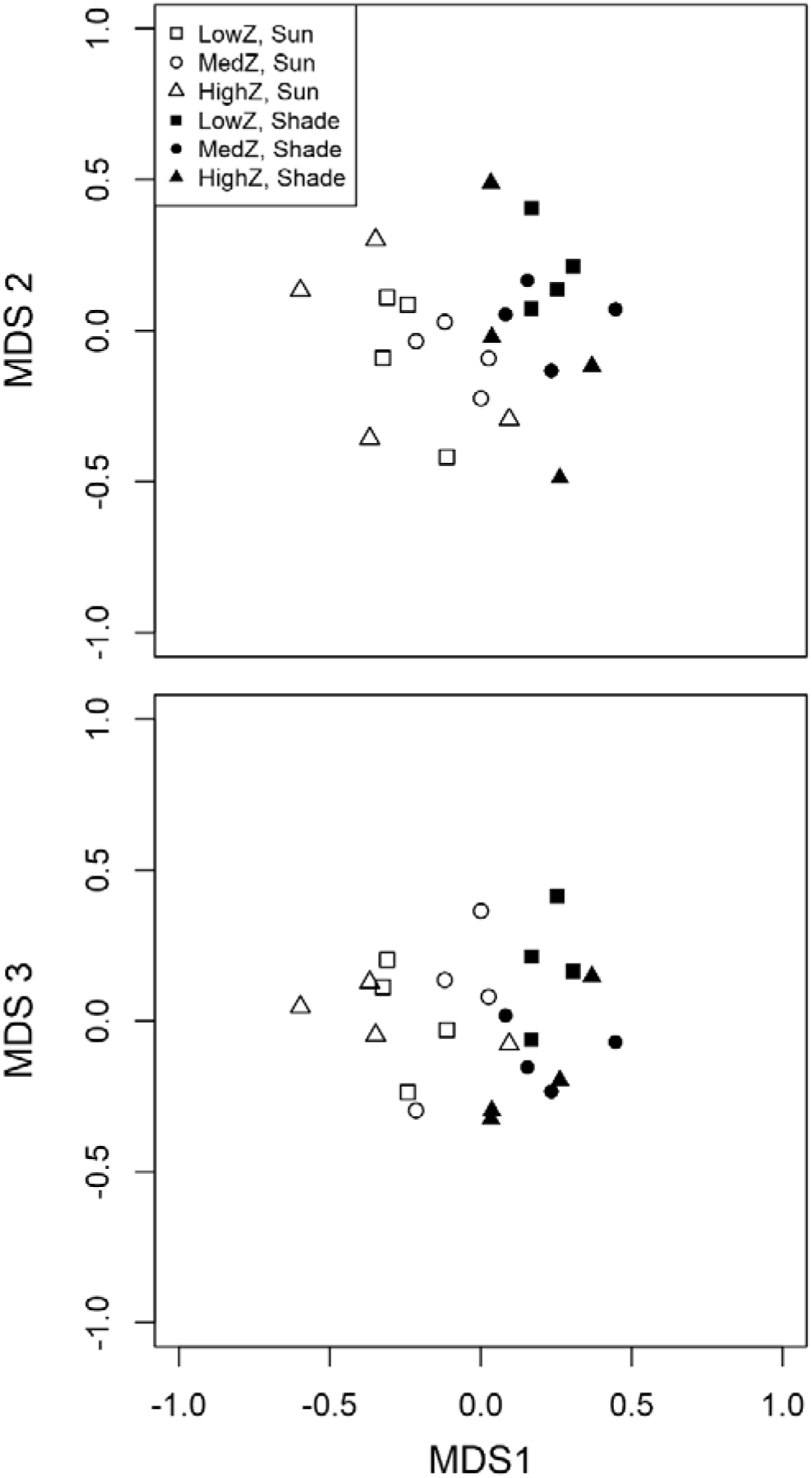
Results from nMDS analysis on phytoplankton community composition with three axes (stress = 0.15). Each point represents one replicate.

Rotifer abundance was not dependent on experimental conditions, but rotifer biomass was significantly affected by zooplankton levels. Rotifer biomass increased with increased zooplankton (Fig. 2; Table 1). Changes in zooplankton treatments accounted for 33% of the variation in rotifer biomass (i.e., *R*^*2*^ _*zoop*_ = 0.33). *Keratella cochlearis* was the dominant rotifer species found in mesocosms and made up > 90% of individuals, but only 39% of biomass due to their small size (Fig. S5). Other rotifers found in mesocosms were *Keratella hiemalis, Brachionus angularis, Asplanchna spp.*, and *Filinia spp*.

Some nutrient levels were significantly different between treatments (Fig. 6; Table 1). SRP and ammonia significantly decreased with light and significantly increased with zooplankton. Light explained 73% of the variation in SRP and 72% of the variation in ammonia, while zooplankton explained 15% of the variation in SRP and 21% in ammonia (Table 1).

**Fig. 6.**
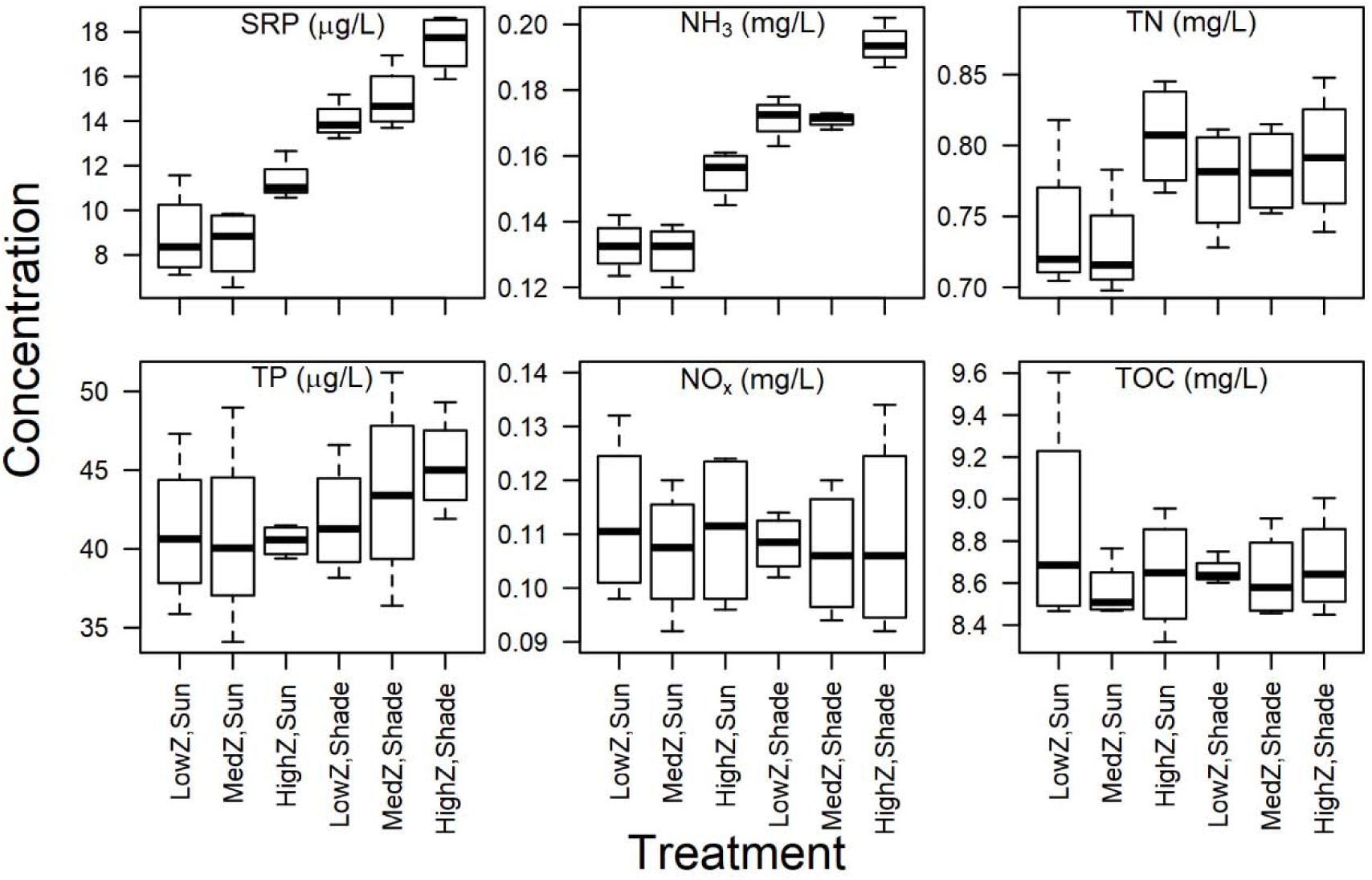
Nutrient results from experimental treatments. “Z” indicates zooplankton level. Center lines indicate medians, boxes indicate first and third quartiles of data, and whiskers indicate minimum and maximum of data.

Zooplankton also significantly increased TN and explained 20% of the variance between treatments (Table 1). Neither light nor zooplankton significantly affected TP, NO_x_, or TOC (Fig. 6; Table 1).

## Discussion

We found that light had a stronger influence on phytoplankton biovolume and community composition than zooplankton grazing in our under-ice carboy experiment. The results support our hypothesis that bottom-up control exceeds top-down control on phytoplankton under ice. Specifically, the greatest contrast in chrysophytes and diatoms occurred when light was manipulated. However, zooplankton grazing had non-negligible effects on both phytoplankton and nutrient concentrations. The significance of zooplankton in our experiment contrasts with the PEG Model assumption that the effects of zooplankton grazing are low during winter (Sommer et al. 1986, 2012). Actively overwintering zooplankton have the potential to impact phytoplankton biovolume under ice, likely through selective feeding and nutrient cycling. Thus, the role of winter zooplankton may be more nuanced than primarily functioning as a standing stock to graze the spring phytoplankton bloom after ice-out as proposed in the “overwintering zooplankton” scenario of the updated PEG Model (Sommer et al. 2012). Changes in phytoplankton community dynamics with zooplankton biomass mediated through light conditions could have direct consequences for which phytoplankton communities are present at ice-out.

Light had a stronger effect on phytoplankton community composition than zooplankton, supporting the hypothesis that light is the main driver of phytoplankton biovolume under ice. The difference was apparent in both nMDS visualizations and ANOVA analysis of specific phytoplankton groups. Changes in community composition contributed to differences in overall phytoplankton biovolume, despite relatively constant total phytoplankton abundance across treatments. In particular, we observed higher proportion of chrysophytes (mostly *Chrysococcus*) and diatoms when more light was available. *Chrysococcus* can be successful under the ice in north temperate lakes (Phillips and Fawley 2002) so *Chrysococcus* as a dominant phytoplankton in Shelburne Pond during winter is not surprising. Although *Chrysococcus* is a known mixotroph that can withstand low light conditions (Olrik 1998), it still responded strongly to high light in our experiment. Diatoms are also often abundant under clear ice with high light transmission similar to our unshaded experimental conditions (Katz et al. 2015). Interestingly, the changes we observed in phytoplankton community composition took place over just two weeks, indicating that highly variable or rapidly changing light environments (e.g., patchy snow, rapid snowfall on top of clear ice, or snow that is abruptly windswept off ice) could have large impacts on phytoplankton community structure over relatively short time scales.

Light treatments significantly altered ammonia and SRP presumably through increased phytoplankton production. NO_x_ decreased in treatments with high phytoplankton levels, but ammonia remained constant across treatments. Many phytoplankton can take up nitrate (the main component of our NO_x_) faster than ammonia, and in some cases, ammonia uptake may inhibit nitrate uptake, although these relationships can be highly variable (Glibert et al. 2016). SRP and nitrate most likely did not limit phytoplankton growth in mesocosms because concentrations of both were higher than concentrations found in Shelburne Pond during summer when phytoplankton production is highest.

Despite discrepancies between expected and actual zooplankton biomass, zooplankton still had a major effect in carboys through consumption of certain phytoplankton groups and alteration of nutrient concentrations. Zooplankton significantly decreased total phytoplankton biovolume, cryptomonad biovolume, and diatom abundance, suggesting that zooplankton selectively grazed larger cryptophytes (specifically *Cryptomonas*) and smaller or non-colonial diatoms. Diatoms and cryptophytes are known food sources for both rotifers and crustacean zooplankton (Mohr and Adrian 2002; Zhou et al. 2011; Tõnno et al. 2016). Furthermore, nutrients showed signals of zooplankton activity through increases in SRP, ammonia, and TN with increased zooplankton. Zooplankton increase ammonia and SRP through excretion (Oliver et al. 2015), whereas higher TN was likely the direct result of higher zooplankton biomass. Although top-down control from zooplankton grazing was minimal in comparison to bottom-up control from light limitation in this experiment, zooplankton have the potential to indirectly affect phytoplankton though mobilization of nutrients during a period when external nutrient inputs are low.

Rotifers unexpectedly increased in biomass as crustacean zooplankton increased. We would expect rotifers to decrease when crustacean zooplankton are abundant because crustacean zooplankton typically outcompete rotifers when phytoplankton resources are limited (Fussmann 1996), and copepods such as *Diacyclops* may directly consume rotifers (Ciros-Pérez et al. 2004). One possibility is that the increased nutrients from adding zooplankton increased primary production that was then consumed by zooplankton and rotifers. However, we cannot evaluate whether nutrient cycling and phytoplankton regeneration rates increased because we only measured standing stock of phytoplankton at the end of experiments. Another potential explanation is that additional rotifers were added with the zooplankton treatments. However, this possibility is unlikely because samples from the 350-µm mesh sieve that were prevented from going into the barrels indicated extremely low rotifer densities of 0.26 individuals/L. Another possibility is that rotifers experienced competitive release if large crustacean zooplankton outcompeted small crustacean zooplankton that may consume phytoplankton in the same size range as rotifers. In this case, having a high prevalence of large crustacean zooplankton would release rotifers from competition. This possibility seems most likely because small-bodied zooplankton were found in lower proportions in treatments with more large-bodied zooplankton.

Mesocosms were effective in maintaining physical parameters such as light and temperature but were less predictable in maintaining zooplankton levels. Water temperatures increased throughout the course of the experiment similarly between shaded and unshaded treatments, so temperature was not an influential factor in differences between treatments. The shade cloth covers maintained differences in light readings between treatments, including during a significant snowfall the night before we began extracting carboys at the end of the experiment (February 8). Zooplankton maintained differences between treatments, but at lower magnitudes than expected. The most likely explanation is that zooplankton experienced mortality as they were collected, sieved, and added to carboys. Alternatively, the highest zooplankton densities may have exceeded the carrying capacity of the carboys and experienced mortality during the experiment.

Mesocosm studies are necessarily limited in their scope to manipulate only the factors under study. In this experiment, we suspended our sealed carboys just below the ice. The limited mixing with the rest of the water column negated the potential for phytoplankton to settle to the bottom of the lake. However, our experiments were conducted during a period when the water was warming before ice-out and was likely in a convective mixing state (Bruesewitz et al. 2015). Convective mixing could re-suspend phytoplankton such as diatoms (Vehmaa and Salonen 2009), and thus, we would not expect phytoplankton to settle out as quickly as they would in a winter stratified state. Additionally, our sealed systems may have relaxed selective pressures for flagellated phytoplankton taxa by limiting their ability to migrate in the water column.

In this experiment, we demonstrated that light is the main driving factor of phytoplankton biovolume and community structure under ice. Variations in light can also lead to significant changes in nutrient cycling. However, the role of zooplankton under ice should not be overlooked. Zooplankton decreased some phytoplankton taxa and altered nutrient concentrations in mesocosms, which suggests that we may miss important contributions of zooplankton in shaping phytoplankton communities and nutrient cycling under ice if we assume that overwintering zooplankton have negligible effects. Furthermore, high prevalence of copepod nauplii suggests that some crustacean zooplankton reproduced under ice. Winter copepod reproduction is often overlooked in temperate lakes, despite its occurrence in multiple systems (this study; Vanderploeg et al. 1992). Additional application of open water experimental techniques to ice-covered ecosystems is an important step in disentangling food web drivers under ice.

The results of this experiment are relevant to understand what may happen with changing winter conditions associated with climate change. As climate change progresses, some regions are expected to experience increased snowpack and other areas are expected to see decreased snowpack (Räisänen 2008). Most regions are also predicted to have earlier snowmelt (Klein et al. 2016) that is potentially more protracted (Musselman et al. 2017). Our study which simulated a change in snow cover of just a few centimeters (Bolsenga and Vanderploeg 1992) for two weeks was enough to significantly alter phytoplankton community structure, indicating that minor events such as rain-on-snow events that melt a layer of snow or slightly altered snowfall totals may have disproportionately large effects on phytoplankton communities. Ice cover duration is expected to shorten in most regions as a result of climate change (Sharma et al. 2019), and while outside the scope of this study, we would expect that shorter ice duration would alter spring mixing regimes (Roberts et al. 2018) and potentially increase zooplankton survival (Preston and Rusak 2010). As indicated in this study, higher zooplankton biomass may increase concentrations of nutrients relevant for phytoplankton growth. Increases in SRP and ammonia could increase nutrient cycling rates if light levels are sufficiently high to sustain phytoplankton growth. The interplay of light transmission and zooplankton grazing and their effects on phytoplankton communities will likely change rapidly with climate change.

## Acknowledgements

We thank Brad Roy, Thomas Hamling, Ben Block, Jessica Griffin, Wilton Burns, Posy Labombard, Taylor Stewart, Brianna Wilkins, Hannah Lister, Natalie Flores, Lisa Izzo, Frances Ianucci, Hannah Lachance, and Brian O’Malley for field work assistance and Jonathan Rickwood for lab assistance. Nicholas Gotelli, Ana Morales-Williams, and Andrew Schroth were instrumental in project design. We also thank the Marsden and Stockwell laboratories for comments on earlier drafts of the manuscript. This work was supported by the University of Vermont Biology Department, the Morse Fund, the UVM Roberto Fabri Fialho Research Award to ARH, the Vermont Space Grant Consortium under NASA Cooperative Agreement NNX15AP86H, and a NASA Earth and Space Science Fellowship to ARH under NASA Cooperative Agreement 80NSSC18K1394 P00001.

