## Supplemental Information for "Under-ice mesocosms reveal the primacy of light but the importance of zooplankton in winter phytoplankton dynamics"

### Supporting Information

**Fig. S1.** Light and water temperatures during the experiment. Light and external temperature were tracked using MK-9 sensors, while internal carboy temperatures were recorded on HOBO loggers.

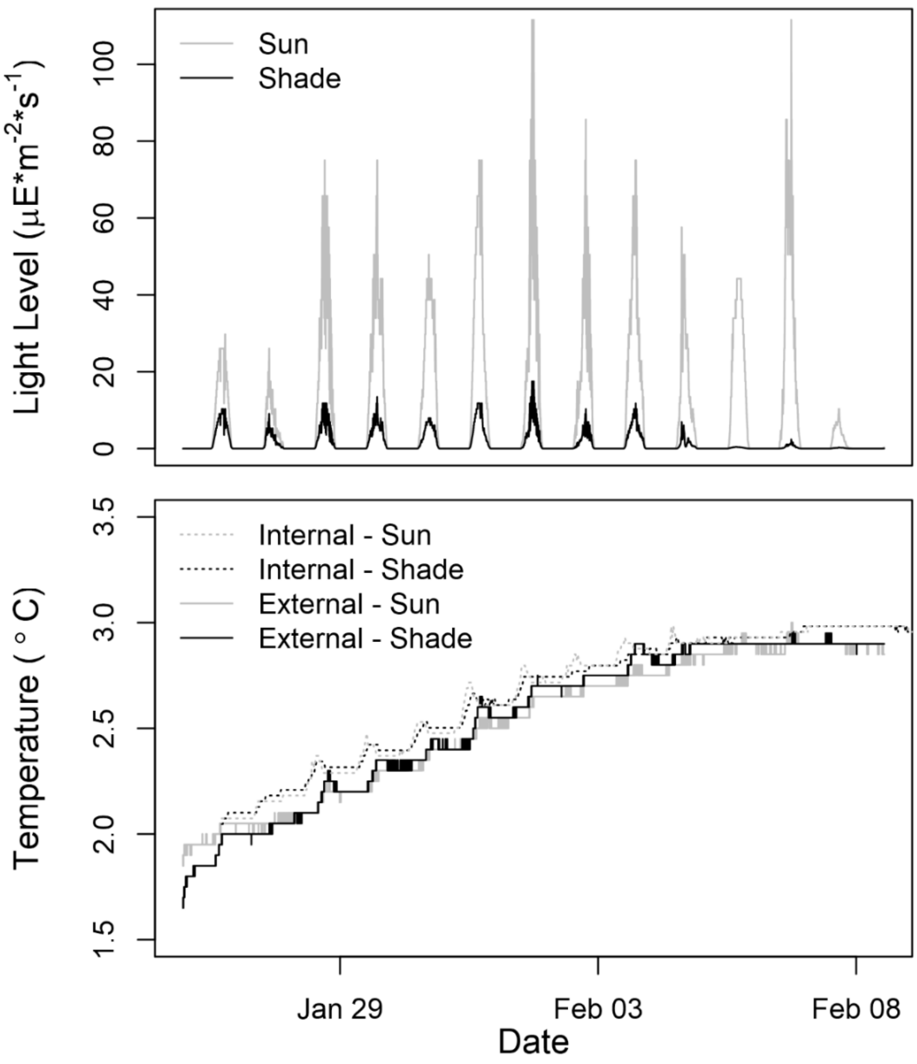

**Fig. S2.** Proportional zooplankton abundance and biomass by species. "Z" indicates zooplankton level.

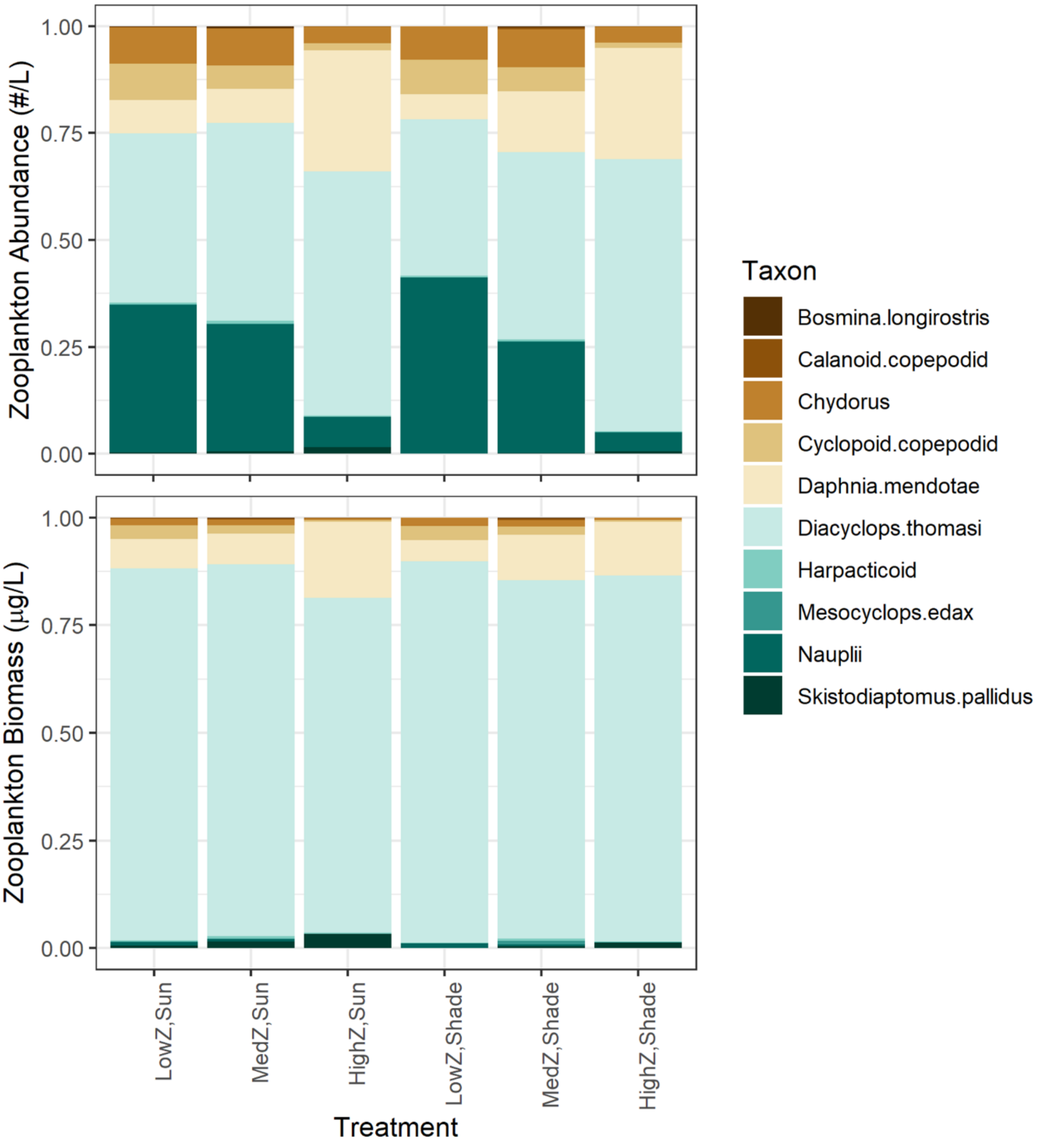

**Fig. S3.** Body size distributions for rotifers and crustacean zooplankton. Note that due to differences in processing protocols and equipment, rotifer and crustacean zooplankton represent different sample sizes. All crustacean zooplankton were measured, whereas the first ten rotifers of each species for each sample were measured. "Z" indicates zooplankton level.

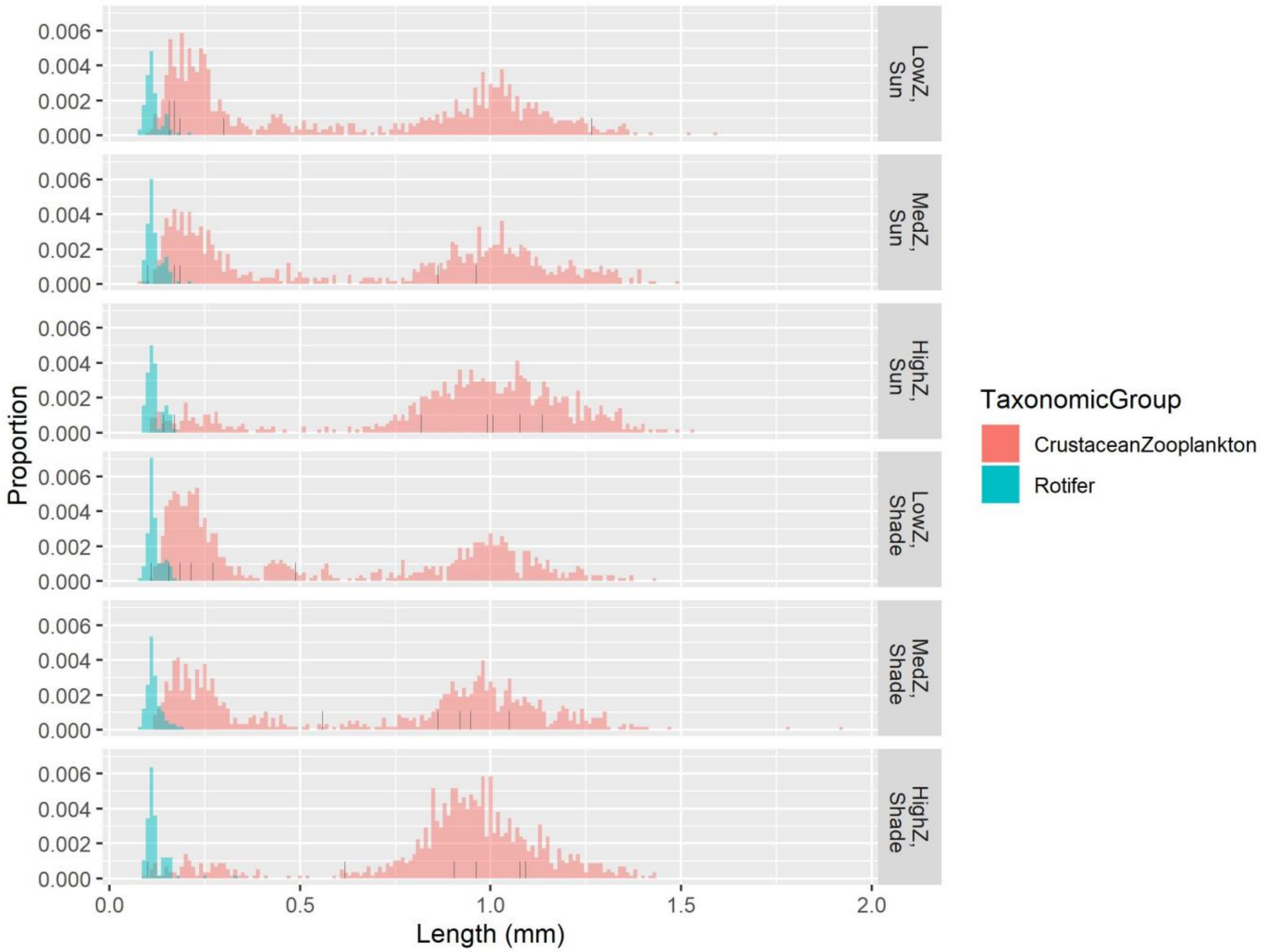

**Fig. S4.** Weights for each taxon contributing to separation in nMDS (Fig. 5). The most positive and most negative weights indicate strong weights on a given axis in opposite directions. Taxa with weights near zero had little impact on separations between phytoplankton communities.

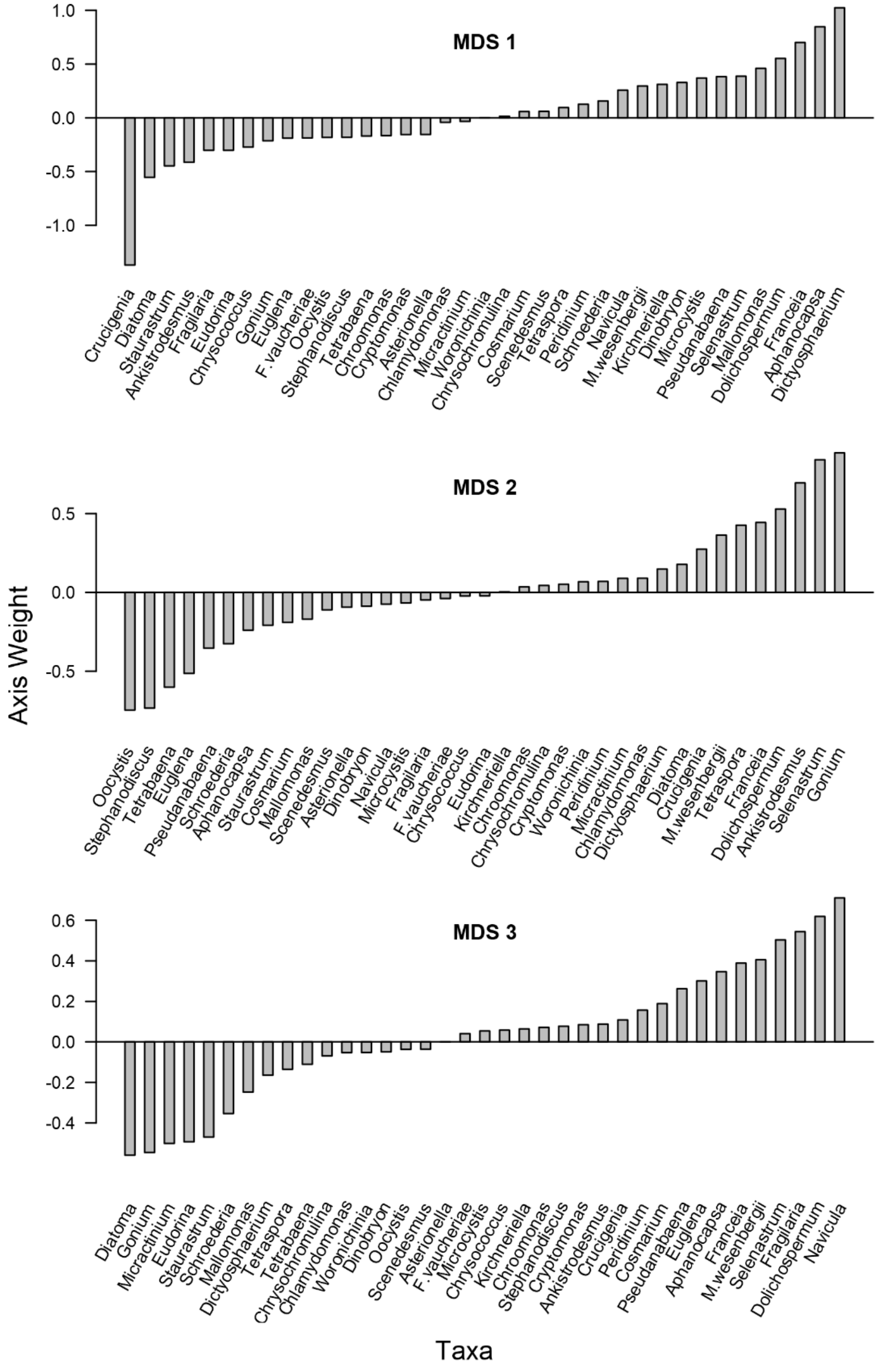

**Fig. S5.** Proportional rotifer abundance and biomass by species. "Z" indicates zooplankton level.

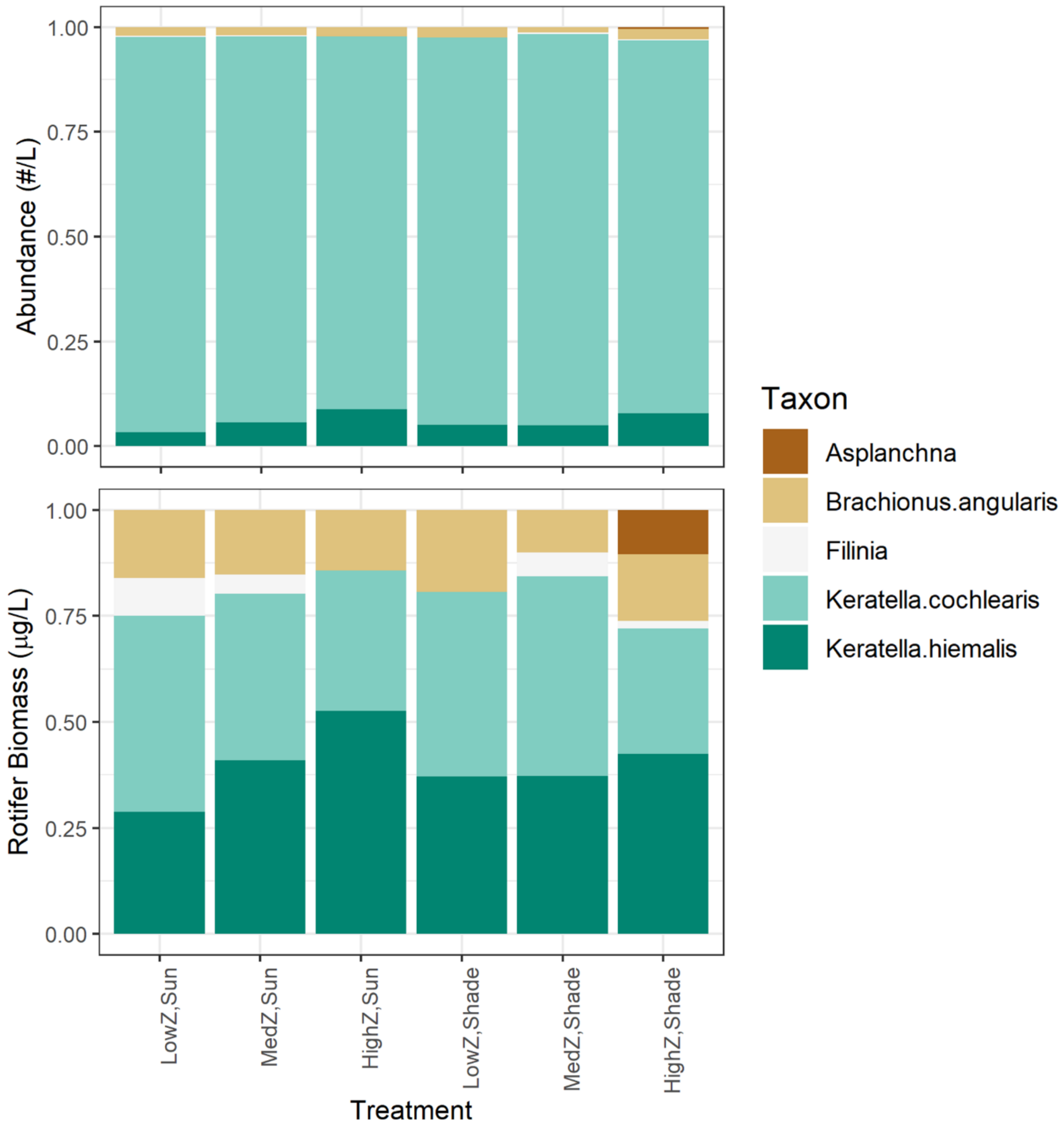
